# Lipid homeostasis is essential for a maximal ER stress response

**DOI:** 10.1101/2022.10.27.513991

**Authors:** Gilberto Garcia, Hanlin Zhang, Sophia Moreno, C. Kimberly Tsui, Brant Michael Webster, Ryo Higuchi-Sanabria, Andrew Dillin

## Abstract

Changes in lipid metabolism are associated with aging and age-related diseases, including proteopathies. The endoplasmic reticulum (ER) is uniquely a major hub for protein and lipid synthesis, making its function essential for both protein and lipid homeostasis. However, it is less clear how lipid metabolism and protein quality may impact each other. Here, we identity *let-767*, a putative hydroxysteroid dehydrogenase, as an essential gene for both lipid and ER protein homeostasis. Knockdown of *let-767* reduces lipid stores, alters ER morphology in a lipid-dependent manner, and also blocks induction of the Unfolded Protein Response of the ER (UPR^ER^). Interestingly, a global reduction in lipogenic pathways restores UPR^ER^ induction in animals with reduced *let-767*. Specifically, we find that supplementation of 3-oxoacyl, the predicted metabolite directly upstream of *let-767*, is sufficient to block induction of the UPR^ER^. This study highlights a novel interaction through which changes in lipid metabolism can alter a cell’s response to protein-induced stress.

## Introduction

The cell must monitor protein and lipid quality to maintain cellular homeostasis. Disruptions in protein folding have been implicated in numerous neurodegenerative diseases, while lipid imbalances result in an increased risk for cardiovascular disease, diabetes, and various cancers [1]–[4]. These two distinct metabolic pathways have been extensively studied independently, yet increasing evidence suggests that examining their relationship to one another may provide novel insights into human health. Individuals with Alzheimer’s disease (AD) and Parkinson’s disease (PD), diseases known to be associated with increased development of abnormal protein aggregates, also show dysregulation of lipid metabolism [5]–[7]. As such, changes in lipid metabolism are now suspected to contribute to the development of these proteopathies through mechanisms that are not fully understood [8]–[10]. The endoplasmic reticulum (ER) is a major metabolic hub, responsible for the synthesis of secreted and integral proteins, as well as a major portion of a cell’s lipids [11]. This close relationship between protein and lipid quality control within the ER is highlighted by the Unfolded Protein Response of the ER (UPR^ER^), a system capable of sensing and responding to both protein misfolding and membrane lipid disequilibrium [12], [13].

In higher eukaryotes, the UPR^ER^ contains three unfolded protein sensors, each with their own independent signaling pathways. While all three sensors are ER-localized transmembrane proteins with a luminal unfolded protein sensing domain, the most conserved of these pathways is the inositol-requiring enzyme-1 (*ire-1*) branch. The IRE-1 luminal domain is bound by the resident ER HSP70 chaperone (*C. elegans* HSP-4) under basal conditions. Upon protein folding stress, the HSP70 chaperone is titrated away to allow for the interaction of the luminal domain with misfolded proteins, leading to oligomerization and activation of IRE-1’s cytosolic RNAse domain. Activated IRE-1 then initiates noncanonical splicing of *xbp-1* mRNA from its *xbp-1u* form to its effective *xbp-1s* variant. The XBP-1s transcription factor is then able to upregulate the cell’s UPR^ER^ target genes to increase protein folding, protein turnover, and lipid metabolism to help ameliorate the stress. [14], [15].

All three of the UPR^ER^ sensors are also able to sense membrane lipid disequilibrium through domains adjacent to their transmembrane helices. These domains activate the UPR^ER^ independent of proteotoxic stress and the luminal sensing domains [16]–[18]. This is unsurprising considering the ER’s critical role in the synthesis of major lipids including membrane lipids, cholesterol, and neutral lipids [19], [20]. Accordingly, the ER actively regulates the cell’s lipid status through changes to lipid synthesis enzymes, transcriptions factors, and interactions with lipid droplets (LDs), organelles tasked with the storage and gatekeeping of surplus lipid stores [21]–[23].

Utilizing the same stress response sensors for protein and membrane lipid stress suggests an interdependent link between lipid and protein homeostasis within the ER. Ectopic activation of the UPR^ER^ results in changes to lipid metabolism, while changes in sphingolipid, lipid saturation, ceramides, and loss of various lipid enzymes activate the UPR^ER^ [16], [18], [24]–[27]. Interestingly, both lipids and LDs have been shown to contribute to proteostasis. A sterol pathway localized to LDs is required for clearing inclusion bodies and LDs themselves are necessary for transport of damaged proteins [28], [29]. Whether other lipid pathways in the ER or on LDs are required for an effective UPR^ER^ response to protein stress has not been fully investigated. Here, we performed a genetic screen of LD associated genes to identify genes whose knockdown affected UPR^ER^ activation in the nematode *C. elegans*. We identified the hydroxysteroid dehydrogenase, *let-767*/HSD17B12, as required for both ER lipid and protein homeostasis. Loss of *let-*767 resulted in severe defects in UPR^ER^ activation, lipid homeostasis, and ER morphology. Defects in ER morphology could be rescued through lipid supplementation, however, the deficiencies in UPR^ER^ activation were not. Instead, knockdown of the upstream regulator of the *let-767* pathway resulted in a significant recovery of the UPR^ER^ activation. Thus, we propose that loss of *let-767* results in accumulation of fatty acid metabolites which leads to detrimental remodeling of the ER membrane and disruption of ER functions in lipid synthesis and UPR^ER^ induction. Our work highlights a unique cellular interaction in which lipid metabolism negatively impacts ER protein homeostasis. Further understanding of how these two systems influence one another may bring new insights into the mechanisms of protein and lipid disorders.

## Results

### LET-767 is a regulator of the UPR^ER^

To identify novel lipid genes that influence the UPR^ER^, we carried out an RNAi screen under conditions of proteotoxic stress. We utilized the *hsp-4* (*C. elegans* Hsp70/BiP) transcriptional GFP reporter (*hsp-4p∷GFP*) to assess the UPR^ER^ induction, and *sec-11* (an ER serine-peptidase) RNAi to generate ER stress and induce the GFP reporter [30]. In combination with the *sec-11* RNAi, we individually knocked down each candidate gene to perform a double RNAi screen **(Fig.1A)**. We focused on proteins associated with LDs instead of general lipid synthesis genes due to the LD’s central role in lipid regulation and their contribution to ER proteostasis [28], [29]. By cross-referencing two independent *C. elegans* LD proteomic datasets, we identified 163 high-confidence LD proteins [31]–[33]. We found that RNA interference of 49 genes led to reduced induction of the UPR^ER^ reporter **(Table S1)**. While a large portion of these genes are annotated as functioning in general translation (*e.g*., ribosomal subunits), potentially affecting global gene expression, a subset of 11 belonged to other functional groups and were therefore more likely to specifically affect the UPR^ER^ **(Fig.S1A).**

**Figure 1.**
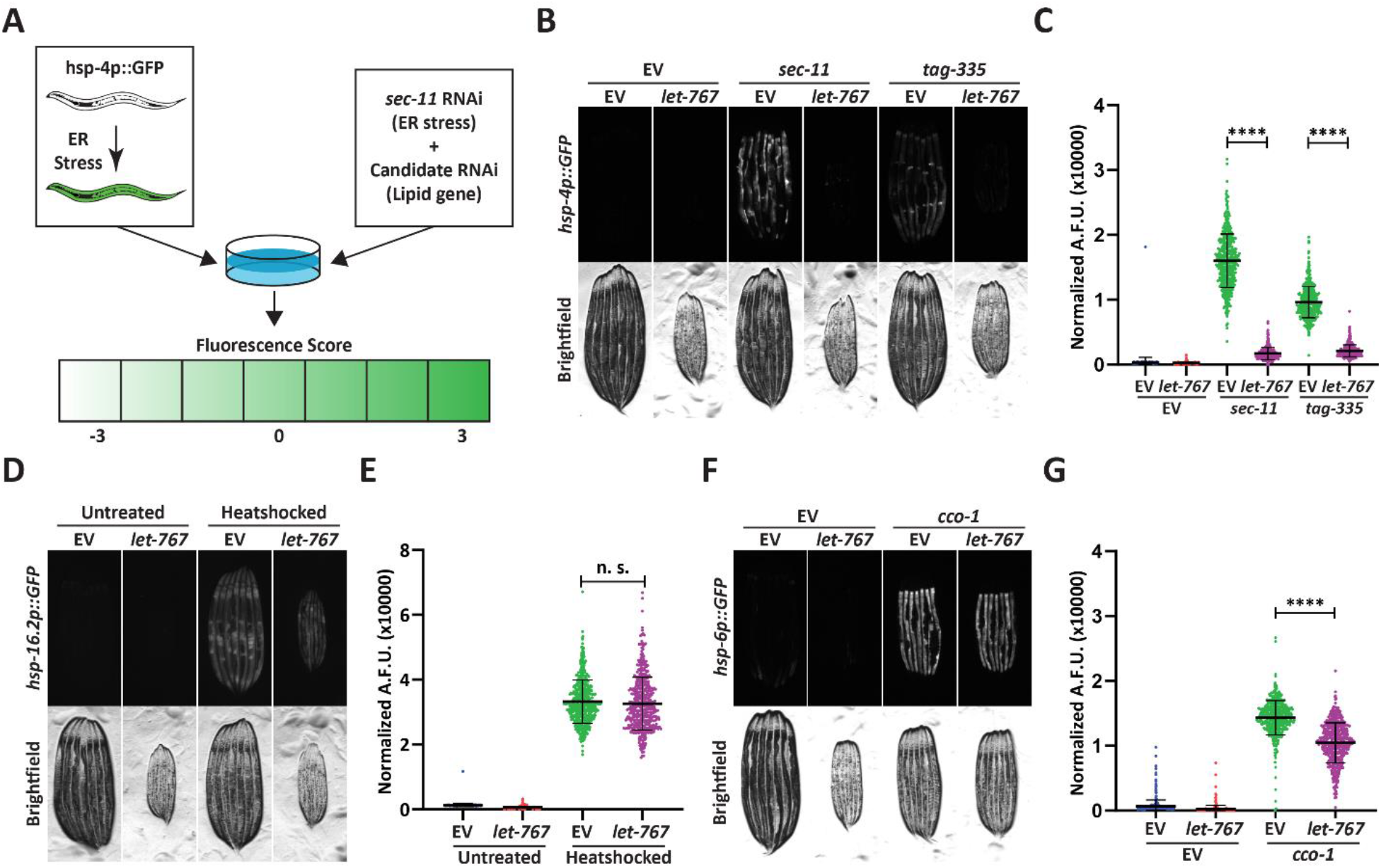
Knockdown of *let-767* specifically suppresses the UPR^ER^. (A) Schematic for screening method used to identify UPR^ER^ modulators from candidate genes. Animals expressing *hsp-4p∷GFP* were grown from L1 on candidate RNAi mixed in a 1:1 Ratio with ER stress inducing *sec-11* RNAi. Animals were then screened at day 1 of adulthood and scored for changes in fluorescence compared to the *sec-11*/Empty Vector (EV) control. (B) Fluorescent micrographs of day 1 adult transgenic animals expressing *hsp-4p∷GFP* grown from L1 on EV, *sec-11*, or *tag-335* RNAi combined in a 1:1 ratio with either EV or *let-767* RNAi to assay effects on UPR^ER^ induction. (C) Quantification of (B) using a BioSorter. Lines represent mean and standard deviation. n = 500. Mann-Whitney test P-value **** < 0.0001. Data is representative of 3 replicates. (D) Fluorescent micrographs of day 1 adult transgenic animals expressing *hsp-16.2p∷GFP* grown from L1 on EV or *let-767* RNAi with or without 2-hour 34°C heat-shock treatment to assay heat shock response. Animals imaged 2 hours after recovery at 20°C. (E) Quantification of (D) using a BioSorter. Lines represent mean and standard deviation. n = 400. Mann-Whitney test n.s.= not significant. Data is representative of 3 replicates. (F) Fluorescent micrographs of day 1 adult transgenic animals expressing *hsp-6p∷GFP*, grown from L1 on EV or *cco-1* RNAi combined in a 1:1 ratio with either EV or *let-767* RNAi to assay effects on UPR^mt^ induction. (G) Quantification of (F) using a BioSorter. Lines represent mean and standard deviation. n = 431. Mann-Whitney test P-value **** < 0.0001. Data is representative of 3 replicates.

We focused on *let-767,* which has been previously characterized to directly affect lipid metabolism [34]–[36]. As a hydroxysteroid dehydrogenase, *let-767* has been implicated in long-chain fatty acid (LCFA) production, monomethyl branched-chain fatty acid (mmBCFA) synthesis, and steroid metabolism. Knockdown of *let-767* suppressed the UPR^ER^ induction caused by RNAi of two different ER genes, *sec-11*, which would impact ER signal peptide cleavage, and *tag-335*, a GDP-mannose pyrophosphorylase, which would reduce protein glycosylation **(Fig. 1B-C)** [37]. Furthermore, qPCR analysis confirmed that *let-767* RNAi also resulted in reduced splicing of *xbp-1* to *xpb-1s* while under stress, suggesting that loss of *let-767* suppresses the UPR^ER^ induction by preventing IRE-1’s splicing of *xbp-1* **(Fig. S1B).** Next, we utilized the transcriptional reporters of the mitochondrial unfolded protein response (UPR^mt^, *hsp-6p∷GFP*) and the heat shock response (HSR, *hsp-16.2∷GFP*) to determine the effect of *let-767* knockdown on other stress responses. We observed that *let-767* RNAi does not suppress the heat shock response and only has minor effects on the mitochondrial stress response in comparison to the highly suppressed UPR^ER^ **(Fig. 1D-G)**. Together, these results show that the function of *let-767* is specifically important for ER function and homeostasis.

To determine whether reduced UPR^ER^ induction was a general phenotype of steroid dehydrogenase knockdown, we performed our double RNAi protocol on the four most closely related (steroid dehydrogenase family) genes found in *C. elegans*, *stdh-1,2,3* and *4*. Knockdown of any of the *stdh* genes had only mild effects on the UPR^ER^ induction, showing that the diminished UPR^ER^ induction phenotype was more specific to *let-767* and not a general phenomenon of decreased steroid dehydrogenase function **(Fig. S1C-D)**. We then tested whether RNAi of other genes implicated in the LCFA and mmBCFA pathways also affected the UPR^ER^ induction [38]–[40]. Of these genes, *acs-1* had the most similar phenotype to *let-767*, while *elo-5* and *hpo-8* had a significant, but less severe effects on the UPR^ER^ induction without stalling development, like *pod-2* **(Fig. S1E-F)**. These results indicate that perturbation of the LCFA/mmBCFA pathways negatively impacts the UPR^ER^, with *let-767* and *acs-1* being the most critical.

### Loss of LET-767 function impacts the UPR^ER^ independent of lipid depletion

Phenotypes caused by a reduction in the LET-767 enzyme could likely result from insufficient production of key metabolites, *i.e.*, mmBCFAs or LCFAs. In such an instance, supplementation of these metabolites would potentially restore the UPR^ER^ induction to its wild type levels. To this end, we supplemented animals with a crude lysate composed of homogenized N2 (wild type) adult animals to provide a complete panel of lipids. We observed that supplementation of lysate was sufficient to rescue the phenotypes of *acs-1* RNAi, including suppression of the UPR^ER^ upon ER stress **(Fig.S2A-B).** However, lysate supplementation of *let-767 RNAi* treated animals resulted in a significant improvement in organismal size, but only a slight improvement in the UPR^ER^ induction **(Fig.2A-B).** This partial rescue suggests that although the animal size is likely the result of insufficient lipids, the suppression of the UPR^ER^ is potentially not due to the absence of key lipids.

**Figure 2.**
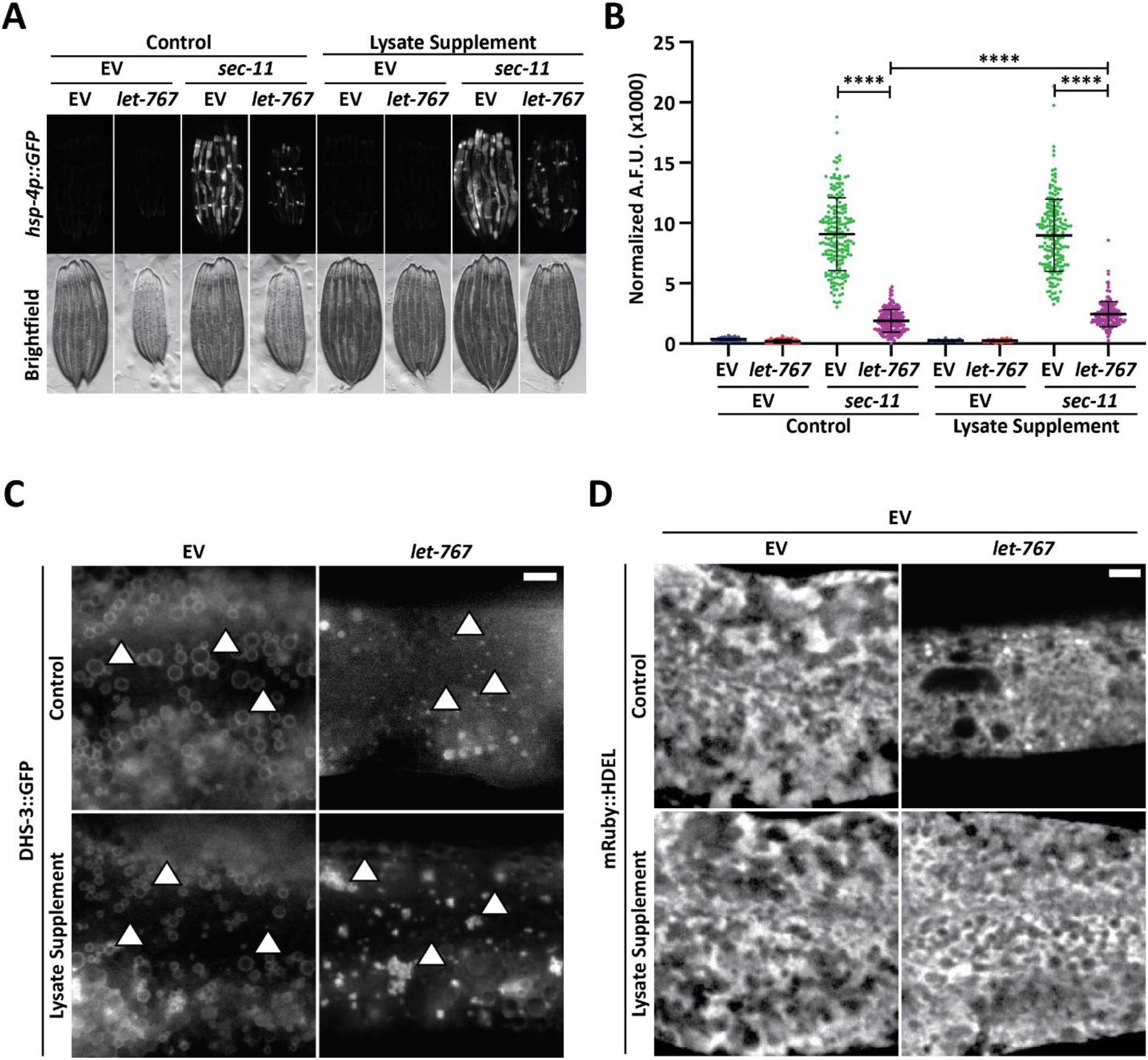
Supplementation of lysate does not restore the UPR^ER^ suppressed by *let-767* RNAi. (A) Fluorescent micrographs of transgenic animals expressing *hsp-4p∷GFP* grown on Empty Vector (EV) or *let-767* RNAi combined in a 1:1 ratio with either EV or *sec-11* RNAi supplemented with vehicle or N2 lysate to assay effects on UPR^ER^ induction. (B) Quantification of (A) using a BioSorter. Lines represent mean and standard deviation. n = 189. Mann-Whitney test P-value **** < 0.0001. Data is representative of 3 replicates. (C) Representative fluorescent micrographs of day 1 adult transgenic animal expressing LD-localized *dhs-3∷GFP*, grown on EV or *let-767* RNAi with or without N2 lysate supplementation to assay LD quality. White arrows denote example lipid droplets. Scale bar, 5μM. (D) Representative fluorescent micrographs of day 1 adult transgenic animal expressing ER lumen-localized *mRuby∷HDEL*, grown on EV or 50% *let-767* RNAi with or without N2 lysate supplementation to assay ER quality. Scale bar, 5μM. Images for organelle markers individually contrasted for clarity.

To investigate whether lysate supplementation had an impact on ER and lipid sub-cellular phenotypes of *let-767* RNAi, we further characterized the impact of *let-767* RNAi and lysate supplementation on the LD and ER morphologies. We found that knockdown of *let-767* caused an extreme reduction in LD size, from large spheres to small points **(Fig.2C)**. *let-767* knockdown also resulted in substantial changes to the ER morphology, practically eliminating the wider sheet-like structures of the ER. However, unlike the UPR^ER^ induction, supplementation of lysate improved the ER morphology to near wild-type appearance and restored the presence of lipid droplets, albeit at considerably lower abundance and size **(Fig.2C-D)**. These observations mark a distinction between the ability of supplementation to rescue morphological and functional phenotypes of the ER caused by *let-767* RNAi. The rescue of animal size and ER morphology phenotypes by lysate supplementation hint that they are a result of insufficient lipids, while the abilities of the ER to induce the UPR^ER^ and expand lipid droplets are potentially not due to the depleted levels of lipids.

Of note, we sought to determine whether the supplementation of only mmBCFAs or LCFAs could rescue the UPR^ER^ induction of animals with reduced *let-767*. We supplemented animals grown on *let-767* and *sec-11* RNAi with exogenous isoC17, an essential mmBCFA, or oleic acid, a LCFA known to be a significant component of the *C. elegans* fatty acid content and to increase lifespan [24], [39], [41]. mmBCFA supplementation was not sufficient to rescue induction of the UPR^ER^ in *let-767* knockdown animals, showing only a slight improvement in the stress response induction **(Fig.S2C-D)**. However, mmBCFA supplementation was sufficient to rescue phenotypes of *acs-1* RNAi, providing evidence that our supplementation effectively delivered the essential mmBCFA and was also sufficient to rescue functional phenotypes of mmBCFA insufficiency **(Fig.S2E-F)**. Similar to isoC17, supplementation of oleic acid showed a very minor improvement in the ER stress response of *let-767* knockdown animals **(Fig.S2G-H)**. These data show that while the UPR^ER^ dysfunction caused by *let-767* knockdown is likely not due to the specific loss of the essential mmBCFA or LCFA (oleic acid), isoC17 is indeed essential to the UPR^ER^ in the absence of *acs-1*.

### Knockdown of lipid biosynthesis pathways restores the UPR^ER^ signaling under *let-767* RNAi

Since supplementation of a WT mixture of lipids was not sufficient to recover the UPR^ER^ induction, but was able to rescue ER morphology, we considered whether *let-767* knockdown could be compromising the ER membrane through unbound LET-767 partners or accumulation of upstream metabolic intermediates. A disrupted membrane could hinder ER membrane protein function for ER stress signaling as well as for lipid droplet and lipid synthesis enzymes, explaining both phenotypes and why restored ER morphology did not coincide with restored UPR^ER^ function. Furthermore, ER stress through *sec-11* RNAi alone caused a reduction in *let-767* transcript levels **(Fig.S1B)**, suggesting that a reduction of *let-767* itself is not directly suppressing the UPR^ER^ induction, but possibly dependent on the levels of other factors. Instead, we hypothesized that disequilibrium within the *let-767* pathway could be the cause of membrane disruption and ultimately the loss of UPR^ER^ induction.

Therefore, we tested whether reducing potentially accumulated upstream metabolites of *let-767* would improve the UPR^ER^ activation. Since the complete enzymatic pathway for *let-767* has yet to be definitively identified and previous works have implicated the enzyme in multiple pathways, we accomplished this by knocking down the ortholog of human SREBP, *sbp-1*, a major transcriptional regulator of numerous lipogenic enzymes and pathways [42]. Analysis of a published RNAseq dataset of *sbp-1* RNAi treated nematodes showed downregulation of numerous lipid synthesis genes, including *let-767*, which we confirmed through qPCR **(Fig.3A)** [43]. We found that animals grown on *let-767* and *sbp-1* RNAi appeared larger and healthier than on *let-767* RNAi alone. More importantly, knockdown of *sbp-1* was able to significantly improve the UPR^ER^ reporter induction in *let-767* knockdown animals under ER stress from *sec-11* RNAi and effectively restore splicing of *xbp-1* to *xbp-1s* **(Fig.3A-C)**. To test whether the improved UPR^ER^ signaling was indeed a functional restoration of the ER stress response, we performed an ER stress survival assay. As expected, animals treated with *let-767* RNAi have a severe defect in survival when exposed to the proteotoxic stress of Tunicamycin, an inhibitor of N-linked glycosylation. Correlating with our observations of the UPR^ER^ reporter, *sbp-1* knockdown rescued the survival of *let-767* RNAi treated animals to the level of *sbp-1* RNAi alone **(Fig.3D-E, Table S2)**. Interestingly, *sbp-1* RNAi itself reduced the induction level of the UPR^ER^ reporter and the survival rate of animals on Tunicamycin, suggesting that while a reduction in the entire *let-767* pathway could indeed rescue the UPR^ER^ suppression caused by *let-767* RNAi, a global reduction in lipid synthesis enzymes limits the maximal induction of the UPR^ER^.

**Figure 3.**
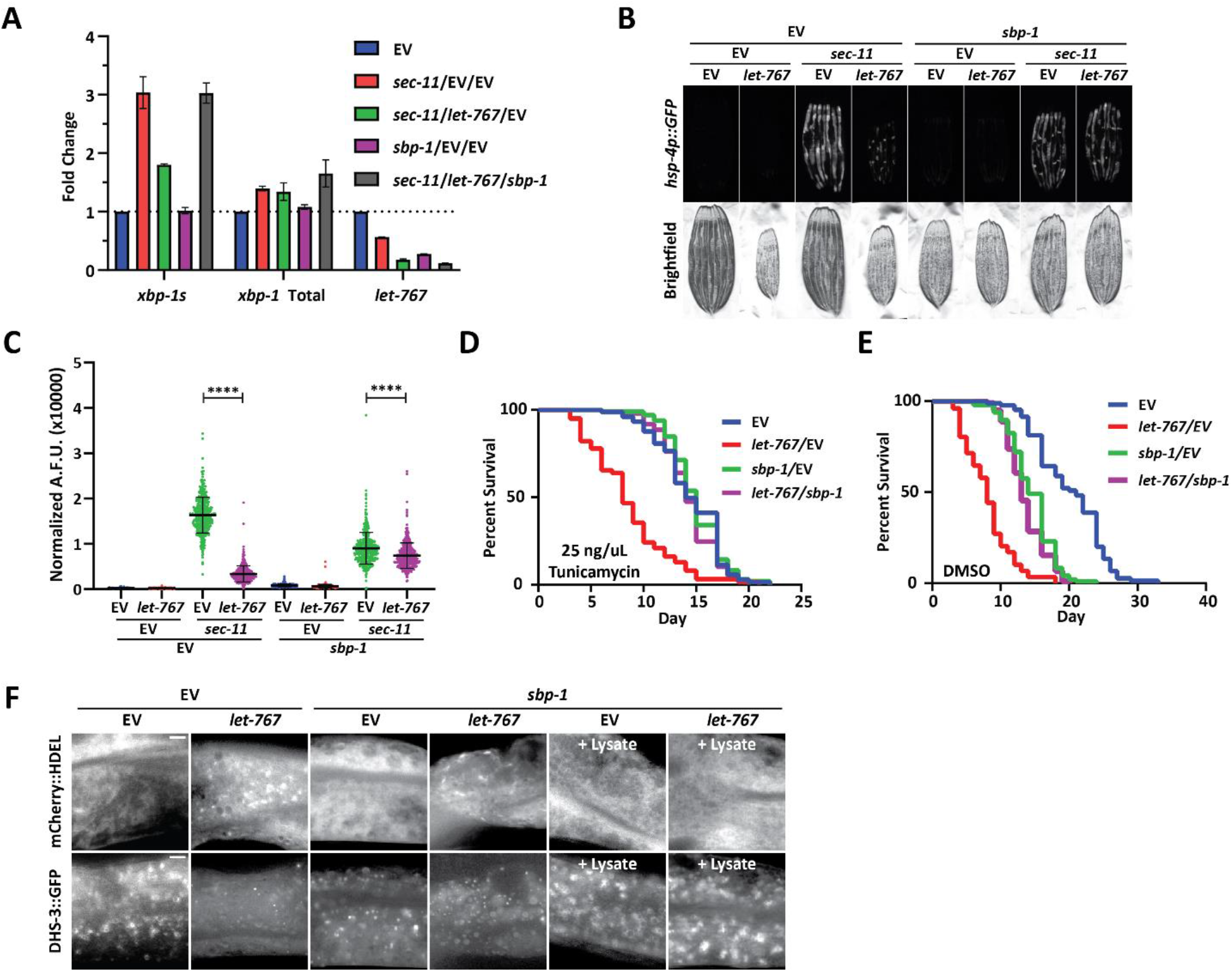
Reduced global lipid synthesis rescues the UPR^ER^ suppression caused by *let-767* RNAi. (A) Quantitative RT-PCR transcript levels of *xbp-1* total, *xbp-1s*, and *let-767* from day 1 adult N2 animals grown from L1 on EV, sec-11 and EV, *let-767* and *sec-11* RNAi combined with either EV or *sbp-1* RNAi in a 1:1:1 ratio. Fold-change compared to EV treated N2 animals. Lines represent standard deviation across 3 biological replicates, each averaged from 2 technical replicates. (B) Fluorescent micrographs of day 1 adult transgenic animals expressing *hsp-4p∷GFP* grown on EV, *let-767*, *sec-11*, and/or *sbp-1* RNAi mixed in a 1:1:1 ratio to assay effects on the UPR^ER^ induction. (C) Quantification of (D) using a BioSorter. Lines represent mean and standard deviation. n = 426. Mann-Whitney test P-value **** < 0.0001. Data is representative of 3 replicates. (D-E) Concurrent survival assays of N2 animals transferred to ER stress conditions of 25 ng/uL Tunicamycin (D) or control DMSO (E) conditions at day 1 of adulthood. Animals continuously grown on EV or *let-767* RNAi combined in 1:1 ratio with EV or *sbp-1* RNAi from L1 synchronization. (F) Representative fluorescent micrographs of day 1 adult transgenic animal expressing ER lumen-localized *mRuby∷HDEL* or LD-localized *dhs-3∷GFP*, grown on EV or *let-767* RNAi mixed in 1:1 ratio with *sbp-1* RNAi with or without N2 lysate supplementation to assay ER and LD quality. Scale bar, 5μM. Images for organelle marker individually contrasted for clarity.

Finally, we determined whether *sbp-1* RNAi would also rescue the defects in LD and ER morphology caused by *let-767* knockdown. Congruent with SREBP’s central role in promoting lipogenesis, we observed a depletion of lipid droplets and a slight perturbation of the ER morphology with *sbp-1* RNAi **(Fig.3F)**. In combination with *let-767* RNAi, *sbp-1* knockdown did not rescue the ER and lipid droplet morphology to WT conditions, highlighting that restoring the UPR^ER^ signaling is not completely dependent on restoring ER lipid levels and morphology. However, in the presence of lysate supplementation, both *sbp-1* RNAi alone or combined with *let-767* RNAi resulted in ER and lipid droplet morphology resembling WT animals. Therefore, a more complete rescue of the phenotypes exhibited by *let-767* knockdown animals could be achieved by a combination of 1) the reduction of the *let-767* pathway through knockdown of *sbp-1* to reduce global lipid synthesis, and 2) exogenous supplementation of the lipids to restore lipids lost by *sbp-1* and *let-767* RNAi.

### *let-767* knockdown impacts *xbp-1* splicing and *xbp-1s* activity

Proper UPR^ER^ signaling is dependent on dimerization of IRE-1 at the ER membrane to splice *xbp-1* mRNA to its active *xbp-1s* isoform. Therefore, one possible mechanism by which loss of *let-767* can impact the UPR^ER^ is by impeding IRE-1 activity through altered membrane dynamics. Indeed, knockdown of *let-767* results in altered ER membrane dynamics where the mobile fraction of an ER transmembrane protein (SPCS-1) is significantly reduced when measured by Fluorescence Recovery After Photobleaching (FRAP) (**Fig.S3A-B**). Supplementation of lysate improved the ER membrane mobility to 62% of WT levels. Comparatively, ER luminal GFP dynamics were less severely impacted by *let-767* RNAi and rescued to a greater extent by lysate supplementation, restoring the mobile fraction of *let-767* RNAi treated animals to 84% of WT levels **(Fig.S3C-D).** While specific changes in dynamics are likely dependent on the individual protein being observed, our results provide evidence that membrane dynamics have been altered when *let-767* is knocked down.

To determine whether the UPR^ER^ was indeed being affected at the ER membrane, we examined the impact of *let-767* knockdown on ectopic UPR^ER^ activation at two different points along the mechanistic pathway: 1) overexpression of *ire-1*, which would ectopically activate the UPR^ER^ at the membrane by constitutively splicing *xbp-1*, and 2) overexpression of the active *xbp-1s* isoform, which would bypass the requirement of IRE-1 splicing at the ER membrane [44]. In creating the necessary strains, we found that intestinal over-expression of the full-length *ire-1a* isoform proved to be lethal. However, overexpression of the *ire-1b* isoform lacking the luminal domain, fused with the mRuby fluorophore (*mRuby∷ire-1b*), was viable and had constitutive activation of the UPR^ER^. To ensure that our fusion protein was responsible for activating the UPR^ER^, we knocked down *ire-1* through an RNAi targeting the N-terminal region only found in *ire-1a*. The *ire-1a* RNAi was able to suppress the UPR^ER^ signaling in WT animals with and without ER stress but did not eliminate the basal UPR^ER^ signal in animals expressing *mRuby∷ire-1b*. Conversely, RNAi targeting common sequences of both *ire-1a* and *ire-1b* suppressed the UPR^ER^ induction in both WT and animals expressing our *mRuby∷ire-1b* construct, even when ER stress was induced by *sec-11* RNAi. **(Fig.S4A-D)**. Therefore, our *mRuby∷ire-1b* construct was sufficient to induce the intestinal UPR^ER^ without the endogenous *ire-1a*. Likewise, we confirmed that overexpression of our intestinal *mRuby∷xbp-1s* construct was able to induce the UPR^ER^ in an *xbp-1 (zc12)* null-mutant background **(Fig.S4E-F)**. Beyond observing that our construct was sufficient to induce the UPR^ER^, we also found that loss of endogenous *xbp-1* resulted in an even higher level of basal UPR^ER^ induction, potentially due to a negative regulatory role for unspliced *xbp-1*, that has been previously suggested [45].

Next, we tested UPR^ER^ induction of our *ire-1* and *xbp-1s* overexpression strains on *let-767* RNAi. *let-767* knockdown reduced the induction of the UPR^ER^ reporter in *mRuby∷ire-1b* animals **(Fig.4A-B)**. However, *let-767* RNAi also reduced the UPR^ER^ reporter induction in the *mRuby∷xbp-1s*, *xbp-1(zc12)* strain **(Fig.4C-D)**. By utilizing the *mRuby∷xbp-1s*, *xbp-1 (zc12)* null-mutant we eliminated the possibility that the reduced induction was due to the negative regulatory effects of unspliced *xbp-1s*. To ensure that the reduced UPR^ER^ induction was not due to altered expression of our *mRuby∷xbp-1s* construct, we performed qPCR for *xbp-1* in animals grown on *let-767* RNAi. We observed a significant drop in the percent of *xbp-1* splicing in animals expressing *mRuby∷ire-1b* when they were grown on *let-767* RNAi, but not in animals expressing *mRuby∷xbp-1s* **(Fig.4E-F)**. However, the reduced UPR^ER^ induction despite the similar levels of *xbp-1s* transcripts between EV and *let-767* RNAi suggests that in addition to affecting *xbp-1* splicing, *let-767* RNAi is likely affecting *xbp-1s* activity downstream of its splicing.

**Figure 4.**
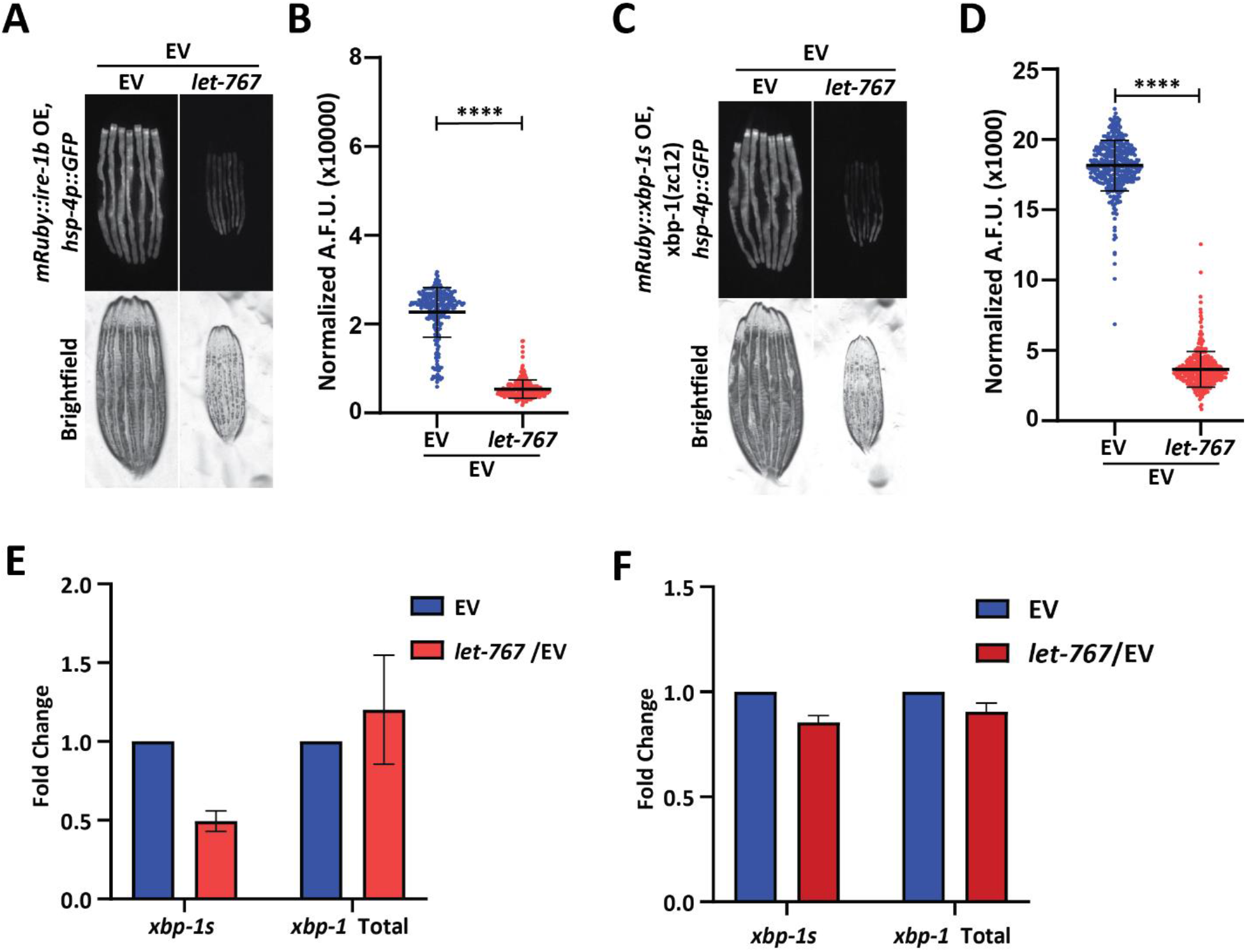
*let-767* knockdown impacts UPR^ER^ induction independent of *xbp-1* splicing. (A) Fluorescent micrographs of day 1 adult transgenic animals expressing *hsp-4p∷GFP* and intestinal *mRuby∷ire-1b* grown on Empty Vector (EV) or 50% *let-767* RNAi to assay the UPR^ER^ induction. (B) Quantification of (A) using a BioSorter. Lines represent mean and standard deviation. n = 290. Mann-Whitney test P-value **** < 0.0001. Data is representative of 3 replicates. (C) Fluorescent micrographs of day 1 adult *xbp-1(zc12)* transgenic animals expressing *hsp-4p∷GFP* and intestinal *mRuby∷xbp-1s* grown on EV or 50% *let-767* RNAi to assay the UPR^ER^ induction. (D) Quantification of (C) using a BioSorter. Lines represent mean and standard deviation. n = 398. Mann-Whitney test P-value **** < 0.0001. Data is representative of 3 replicates. (E) Quantitative RT-PCR transcript levels of *xbp-1s* and total *xbp-1* from day 1 adult *mRuby∷ire-1b* animals grown from L1 on *let-767* RNAi mixed 1:1 with EV. Fold-change compared to EV treated animals. Error bars indicate ± standard deviation across 3 biological replicates, each averaged from 2 technical replicates. (F) Quantitative RT-PCR transcript levels of *xbp-1s* and total *xbp-1* from day 1 adult *mRuby∷xbp-1s* animals grown from L1 on *let-767* RNAi mixed 1:1 with EV. Fold-change compared to EV treated animals. Error bars indicate ± standard deviation across 2 biological replicates, each averaged from 3 technical replicates.

### The *let-767* upstream metabolite, 3-oxostearic acid, reduces induction of the UPR^ER^

To identify a node within the *let-767* pathway that might be responsible for the disruptive metabolite or protein, we utilized mammalian cell culture to probe the effect of specific metabolites in the *let-767* pathway on ER homeostasis. This would allow us to saturate cellular exposure to lipid intermediates and avoid the possibility of bacteria processing the intermediates prior to digestion by *C. elegans* animals. Similar to *let-767* in *C. elegans*, the 3-ketoacyl reductase in mice, HSD17B12, was found to be essential for development in mice and knockout of the gene in adult mice resulted in reduced body weight, reduced lipid content, and caused liver toxicity hypothesized to be from accumulation of toxic intermediates [46], [47]. 3-ketoacyl reductases perform the second step in fatty acid synthesis/elongation, metabolizing 3-oxoacyl-CoA to 3-hydroxyacyl-CoA **(Fig.5A)** [48]. While the fatty acid elongation pathway has not been shown to affect the UPR^ER^, use of the fatty acid synthase inhibitor, Cerulenin, has been shown to increase levels of XBP1s with reduced transcriptional activity due to changes in palmitoylation [49]. From these studies, we hypothesized that the accumulation of the metabolites upstream of LET-767 could be reducing the transcriptional activity of XBP-1S. To this end, we tested whether an upstream metabolite of LET-767 was sufficient to reduce UPR^ER^ induction in huh7 cells containing a 5x unfolded protein response element (UPRE) GFP reporter and EIF2A promoter driving mCherry. As fatty acid elongation extends fatty acids beyond the 16 carbons synthesized by fatty acid synthase, we supplemented our reporter line with the 18 carbon fatty acid metabolites upstream and downstream of the 3-ketoacyl reductase in combination with Tunicamycin to induce ER stress **(Fig.5B-C)**. We observed that the upstream metabolite, 3-oxostearic acid, reduced the normalized 5xUPRE reporter induction. In comparison, the downstream metabolites, 3-hydroxystearic acid and stearic acid, did not reduce the normalized 5xUPRE reporter signal. Instead, stearic acid caused a slight induction of the reporter signal, in agreement with studies showing that saturated lipids induce the UPR^ER^ [50]. This indicated that the 5xUPRE was specifically sensitive to the 3-oxo metabolite upstream of the *let-767* pathway and that this interaction between lipid metabolism and the UPR^ER^ is likely conserved in mammalian cells.

**Figure 5.**
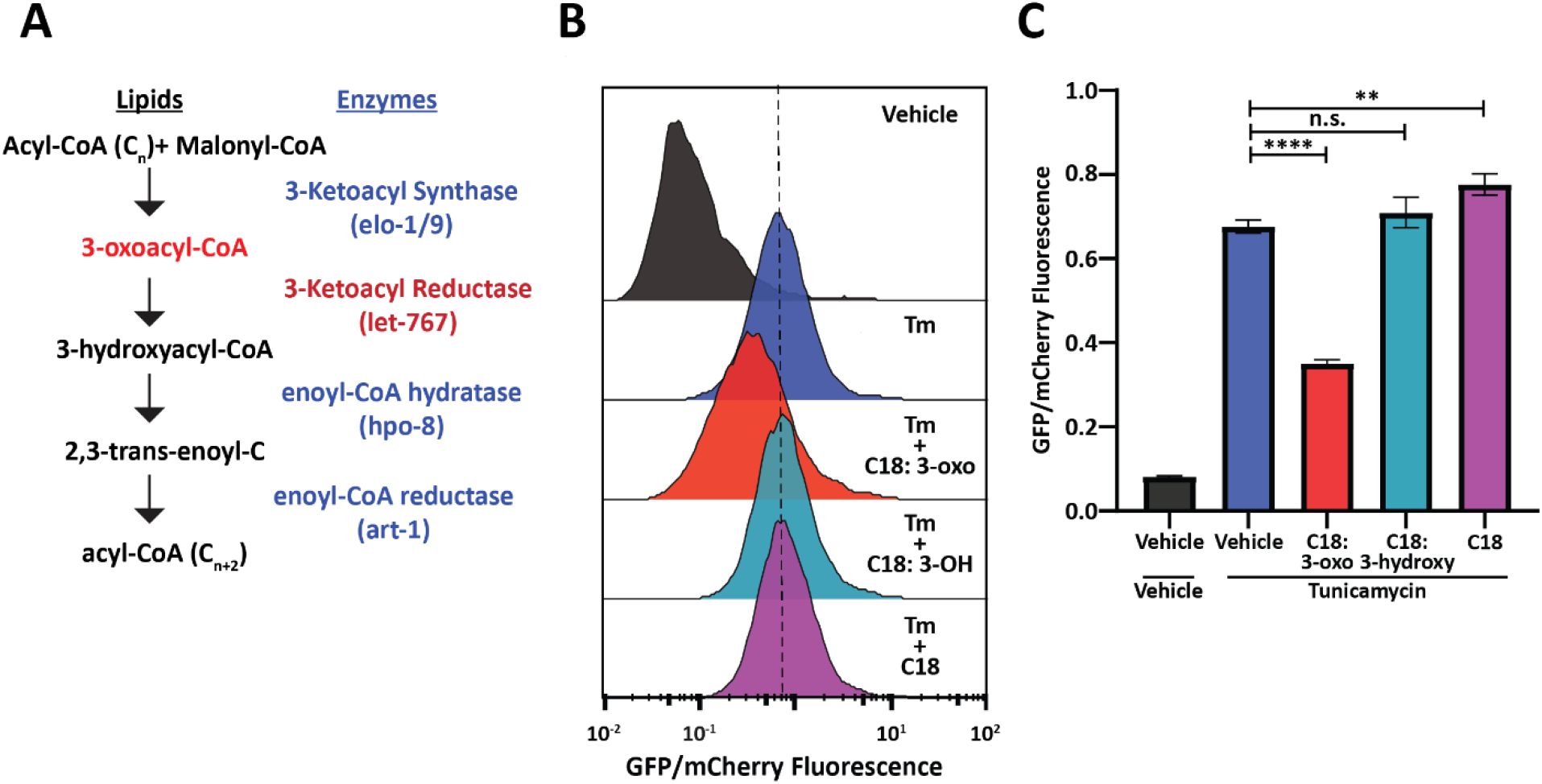
Fatty acid intermediate, 3-oxoacyl, is sufficient to reduce UPR^ER^ induction. (A) Fatty acid elongation pathway displaying intermediate lipid metabolites and annotated *C. elegans* genes. (B) Flow cytometry measurement histogram of huh7 UPRE reporter fluorescence normalized to EIF2A promoter driving mCherry. Cells were treated for 18 hours with Tunicamycin and vehicle, 50 uM Cerulenin, 50 uM 3-oxostearic acid, 50 uM stearic acid, or 50 uM 3-hydroxystearic acid. Data is representative of 2 biological replicates. (C) Median bar graph of (H). Error bars indicate ± standard deviation across 3 technical replicates.

## Discussion

Cells must monitor and maintain both lipid and protein homeostasis to preserve cellular function. The ER is uniquely a major site of both protein and lipid synthesis. Here, we performed a genetic screen to identify lipid regulators that impact the ER response to protein stress and found *let-767* to be necessary for proper animal size, organelle morphology, neutral lipid accumulation, and induction of the UPR^ER^. Under WT conditions, the *let-767*/HSD17B12 pathway elongates fatty acids that are then used to produce other lipids, such as triglycerides stored in lipid droplets **(Fig.6A)**. The UPR^ER^ is also able to respond to ER stress by inducing XBP-1s activity. When the *let-767* is knocked down, the pathway is unable to elongate fatty acids and accumulates the upstream metabolite, 3-oxoacyl-CoA, and possibly other metabolic intermediates that disrupt the ER membrane quality **(Fig.6B)**. Cells are unable to synthesize downstream lipids for lipid droplets and either through disruption of the membrane or direct interaction with 3-oxoacyl-CoA, *xbp-1* splicing and the activity of spliced *xbp-1s* is significantly reduced in the presence of ER stress. While unfolded proteins and lipid disequilibrium are known to independently induce the UPR^ER^, our novel finding demonstrates that changes in lipid pathways could significantly impact a cell’s ability to respond to protein stress and brings to light a potential mechanism through which lipid disequilibrium might facilitate the progression of proteopathic diseases.

**Figure 6.**
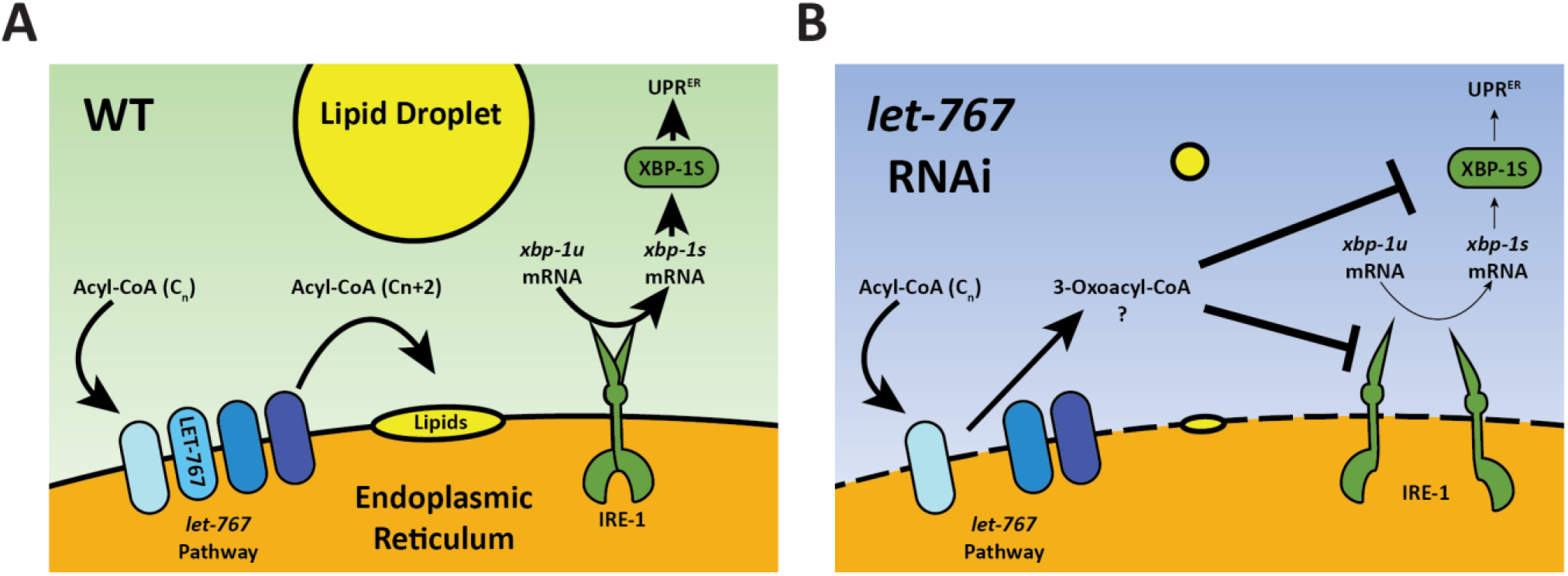
Model for *let-767* RNAi blocking UPR^ER^ induction. (A) Under WT condition, acyl-CoA metabolites are elongated by the *let-767*/HSD17B12 pathway and utilized at the membrane to synthesize other lipids such as neutral lipids stored in lipid droplets. IRE-1 responds to ER stress by splicing *xbp-1u* to *xbp-1s*, which is then translated and able to induce expression of the UPR^ER^ target genes (B) Knockdown of *let-767* results in dysequilibrium of *let-767*/HS17B12 pathway, leading to accumulation of intermediates such as 3-oxoacyl-CoA and reduced lipid production. Intermediate metabolites disrupt membrane quality (dashed line) and negatively affect induction of the UPR^ER^ by reducing splicing of *xbp-1s* (smaller arrows) by IRE-1 and reducing the function of *xbp-1s* post-splicing (smaller arrows).

LET-767 has been characterized as a hydroxysteroid dehydrogenase with evidence as both a steroid modifying enzyme and a 3-ketoacyl reductase located on the ER, consistent with mammalian HSD17B3 and HSD17B12 [34], [35]. LET-767 has also been implicated as a requirement for branched-chain and long-chain fatty acid production, however, its linear metabolic pathway has not been thoroughly investigated so its exact lipid products or protein interactors are yet to be identified. Through our depletion of mmBCFA and LCFA pathway genes, we find that both pathways are essential to having a maximal ER unfolded protein response to protein stress. While individual supplementation of mmBCFAs or LCFAs did not rescue the *let-767* RNAi phenotypes, supplementation of a more complex lipid mixture, crude worm lysate, was able to significantly rescue organelle morphology, size, and reproduction phenotypes. This suggests that loss of *let-767* has a more global effect on lipid production rather than affecting a single lipid species. A straightforward explanation for its requirement in global lipid equilibrium and the UPR^ER^ could be that the *let-767* pathway is crucial for an elemental component of the ER, such as production of fundamental lipids required for ER membrane quality.

We would then expect that *let-767* levels would be critical for ER functions, including UPR^ER^ signaling. Interestingly, ER stress results in reduced *let-767* transcript levels. Furthermore, the reduced UPR^ER^ signaling caused by *let-767* RNAi was significantly rescued by knockdown of the upstream transcription factor, *sbp-1*, which also reduced *let-767* transcript levels and lipid stores. A possible interpretation of these results is that an alternative pathway to produce the essential lipids is upregulated under ER stress or *sbp-1* RNAi, but with the existence of an alternative pathway, the UPR^ER^ would not be suppressed by knockdown of *let-767*. Instead, we propose that the *let-767* RNAi phenotypes are caused by disequilibrium of the *let-767* pathway. By reducing a single part of the *let-767* pathway, intermediate metabolites could interfere with ER membrane dynamics, interactions, and functions. As the UPR^ER^ sensors and numerous lipogenic enzymes reside on the ER membrane, a disrupted ER membrane would explain why knockdown of *let-767* and other lipid genes affects the capacity of the UPR^ER^ induction. While knockdown of *sbp-1* also reduced the level of UPR^ER^ induction, the reduction of numerous lipogenic pathways, including the entire *let-767* pathway, was able to improve the UPR^ER^ function of animals on *let-767* RNAi, potentially by prevent the lipid disequilibrium within the *let-767* pathway.

To test whether the ER membrane disorganization was the source of UPR^ER^ dysfunction, we aimed to bypass the splicing of *xbp-1* by IRE-1 at the ER membrane by overexpressing the already spliced isoform, *xbp-1s*. However, *let-767* RNAi still caused a reduction in the ER stress response. The reduced UPR^ER^ induction in *xbp-1s* overexpressing animals points to an additional mechanism downstream of splicing for the muted UPR^ER^. XBP1u mRNA has been shown to localize to the ER membrane to facilitate splicing to XBP1s upon ER stress [53], [54]. Whether *xbp-1* mRNA localization to the ER membrane is required for further regulation beyond splicing is unknown. If the ER membrane is indeed disrupted by *let-767* RNAi, localization of *xbp-1s* mRNA to the ER membrane would likely be impacted. While splicing is a major point of regulation for *xbp-1*, previous studies have found other potential nodes of regulation for *xbp-1* [49], [51], [52]. Accumulation of the metabolite upstream of LET-767 could be disrupting the membrane in addition to other effects such as reducing fatty acid synthase activity or protein palmitoylation. Through supplementation of fatty acid elongation intermediates to human huh7 hepatocytes, we provide evidence that increased levels of the metabolite upstream of a 3-ketoacyl reductase, 3-oxoacyl, is sufficient to reduce the ER stress response to a similar level as the fatty acid synthase inhibitor, Cerulenin [49]. Whether the 3-ketoacyl metabolite impacts fatty acid synthase activity or protein palmitoylation requires further investigation.

## Materials and methods

### Nematode Strains

N2 Bristol, LIU1 (ldrIs[dhs-3p∷dhs-3∷GFP]), SJ4005 (zcIs4[hsp-4p∷GFP]), SJ17 (xbp-1(zc12) III; zcIs4 [hsp-4p:GFP] V), SJ4100 (zcIs13[hsp-6p∷GFP]), CL2070 (dvIs70[hsp-16.2p∷GFP]), VS25 (hjIs[vha-6p∷GFP∷C34B2.10(SP12) + unc-119(+)]),EG6703 (unc-119(ed3); cxTi10816; oxEx1582[eft-3p∷GFP + Cbr-unc-119]) strains were obtained from the Caenorhabditis Genetics Center (CGC). AGD2192 (uthSi60[vha-6p∷ER-signal-sequence∷mRuby∷HDEL∷unc-54 UTR, cb-unc-119(+)] I; unc-119(ed3) III) [25]. Transgenic strains created for this study were generated from EG6703 via the MosSCI method [55] or through crossing strains.

Transgenic strains created:

AGD2424 (unc-119(ed3) III; uthSi65[vha-6p∷ERss∷mRuby∷ire-1a (344-967aa)∷unc-54 3’UTR cb-unc-119(+)] IV)

AGD2425 (uthSi65[vha-6p∷ERss∷mRuby∷ire-1a (344-967aa)∷unc-54 3’UTR cb-unc-119(+)] IV; zcls4[hsp-4p∷GFP] V)

AGD2012 (unc-119(ed3) III; uthSi71[vha-6p∷mRuby∷xbp-1s∷unc-54 3’UTR cb-unc-119(+)] IV)

AGD2735 (uthSi71[vha-6p∷mRuby∷xbp-1s∷unc-54 3’UTR cb-unc-119(+)] IV; zcls4[hsp-4p∷GFP] V)

AGD2996 (xbp-1(zc12) III; uthSi71[vha-6p∷mRuby∷xbp-1s∷unc-54 3’UTR cb-unc-119(+)] IV; zcls4[hsp-4p∷GFP] V)

### Worm growth and maintenance

All worms were maintained at 20°C on NGM agar plates seeded with OP50 E. coli bacteria. Prior to experiments, worms were bleach synchronized as described in [55], followed by overnight L1 arrest in M9 buffer (22 mM KH2PO4 monobasic, 42.3 mM Na2HPO4, 85.6 mM NaCl, 1 mM MgSO4) at 20°C. For RNAi experiments, arrested L1s were plated on 1μm IPTG, 100 μg/mL Carbenicillin NGM agar plates maintained at 20°C and seeded with RNAi bacteria grown in LB+ 100 μg/mL Carbenicillin

### Fluorescent microscopy

Transcriptional reporter strains were imaged using a Leica DFC3000 G camera mounted on a Leica M205 FA microscope. Worms were grown to day 1 of adulthood at 20°C, hand-picked, and immobilized with 100 mM Sodium Azide M9 buffer on NGM agar plates. Raw images were cropped, and contrast matched using ImageJ software.

For the initial screen of hsp-4p∷GFP reporter animals, fluorescence was scored by eye using the following criteria: 2 = increased fluorescence, 1 = possible increase in fluorescence, 0 = no change, −0.5 = small regions of dimmer fluorescence, −1= small regions of complete loss of fluorescence, −1.5 = globally dimmer fluorescence and some regions of no fluorescence, −2 = global loss of fluorescence and small regions of dim fluorescence, −2.5 = global loss of fluorescence except for regions within spermatheca, −3 = complete loss of fluorescence.

High-magnification fluorescent images were acquired with a Leica DFC9000 GT camera mounted on a Leica DM6000 B microscope. Day 1 worms were picked onto 10 mM Sodium Azide M9 buffer on slides and imaged within 15 minutes. Raw images were cropped and independently contrast optimized for clarity using ImageJ software.

Confocal images were acquired using a 3i Marianas spinning-disc confocal platform. Day 1 worms were picked onto 10 mM Sodium Azide M9 buffer on slides and imaged within 15 minutes. Raw images were cropped and independently contrast optimized for clarity using ImageJ software.

### Biosorter Analysis

Transcriptional reporter strains were grown to day 1 of adulthood at 20°C. Animals were collected into a 15mL conical tube with M9 buffer and allowed to settle at the bottom before supernatant was aspirated. Animals were resuspended in M9 buffer and analyzed using a Union Biometrica COPAS Biosorter (P/N: 350-5000-000) as described in [37]. Animals which saturated the signal capacity of 65532 or were outliers in both animal size parameters (*i.e.*, Extinction and Time Of Flight) were censored. Mann-Whitney statistical tests were performed on fluorescence normalized to animal extinction using GraphPad Prism software.

### qPCR

Animals grown on EV or RNAi bacteria were collected at day 1 of adulthood using M9 and washed 3x. M9 was aspirated and trizol added before 3 cycles of freeze/thaw in liquid nitrogen. Chloroform was then added at a ratio of 1:5 (chloroform:trizol). Separation of RNA was performed through centrifugation in gel phase-lock tubes. The RNA aqueous phase was transferred into new tubes containing isopropanol. RNAi was purified using the QIAGEN RNeasy Mini Kit (74106) according to the manufacturer’s instructions. cDNA was synthesized using 2 μg of RNA and the QIAGEN QuantiTect Reverse Transcriptase kit (205314) according to the manufacturer’s instructions. qPCR was performed using SYBR-green. Analysis was performed for each biological replicate using the Delta-Delta CT method with *pmp-3*, *cdc-42*, and *Y45F10D.4* as housekeeping genes [56].

### Lysate supplementation

Crude lysate was obtained from N2 worms grown on 40x concentrated EV bacteria at 20°C. ~120,000 day 1 adult worms were collected with M9 and washed 6x with M9 buffer. Animals were then homogenized 20x in 3 mL of M9 buffer using an ice-cold 15 mL Dura-Grind Stainless Steel Dounce Tissue Grinder (VWR, 62400-686). Crude lysate was transferred to 1.5 mL tubes and frozen in liquid N2.

Supplementation experiments were prepared by mixing 4x concentrated RNAi bacteria with crude lysate at a 2:1 ratio, respectively. The lysate mixture was plated on RNAi plates and allowed to dry. Dried plates were then UV irradiated without lids for 9 minutes in an ultraviolet crosslinker (UVP, CL-1000) before plating L1 arrested worms on the plate.

### Lipid supplementation

Lipid supplementation experiments were prepared by inoculating cultures (LB + 100 μg/mL Carbenicillin) with RNAi bacteria or empty vector and then adding the lipids to the specified concentration or equal volumes of ethanol. The cultures were allowed to grow overnight to saturation and then concentrated to 4x before being plated on RNAi plates. The bacteria was allowed to dry overnight and then UV irradiated without lids for 9 minutes in an ultraviolet crosslinker (UVP, CL-1000) before plating L1 arrested worms on the plate.

### Tunicamycin survival assay

Tunicamycin survival assays were conducted on NGM agar plates containing 25ng/uL of tunicamycin in DMSO, or equal volume of DMSO with specified RNAi bacteria at 20°C. Animals were moved daily for 4-7 days to new RNAi plates until progeny were no longer observed. Worms with protruding intestines, bagging phenotypes, or other forms of injury were scored as censored and not counted as part of the analysis. For combined RNAi lifespans, saturated cultures were mix 1:1 by volume.

### FRAP analysis

FRAP 4D images were acquired using a 3i Marianas spinning-disc confocal platform. Photobleaching of a 10 μm x 10 μm region within a 4 μm Z-stack was performed using a 488 nm laser for 1-2 ms. Raw images were processed with FIJI [57] into sum Z-projections and aligned using the “RigidBody” setting of the StackReg ImageJ plugin [58]. FRAP analysis was then performed with the FRAP Profiler ImageJ plugin [59] by selecting the 10 μm x 10 μm photobleached region as region 1 and the entire fluorescent area as region 2.

### Cell Culture

Cells were grown in DMEM media (11995, Thermo Fisher) supplemented with 2 mM GlutaMAX (35050, Thermo Fisher), 10% FBS (VWR), Non-Essential Amino Acids (100X, 11140, Thermo Fisher), and Penicillin-Streptomycin (100X, 15070, Thermo Fisher) in 5% CO2 at 37 °C.

### Generation of Huh7 cell lines

To generate a UPRER transcriptional reporter, we designed a lentiviral vector that encodes sequence for 5x UPR response element upstream of a minimal cFos promoter, driving sfGFP [60]. The sfGFP is fused to a PEST sequence for tighter regulation of the reporter [61], [62]. The vector also allows for constitutive mCherry expression for normalization of the sfGFP signal, and neomycin resistant gene for selection. The transcriptional reporter was then transduced through lentivirus into Cas9-expressing Huh7 cells and selected for using G418 at 800 ug/mL.

### Cell Culture Supplementation Experiments

Cells were plated onto 10cm plates and allowed to grow overnight. Lipids, tunicamycin, and/or vehicle were directly added to media. Cells were harvested for flow cytometry 16-18 hours after addition of supplements.

### Flow Cytometry

Cells were trypsinized on ice and resuspended in cold FACS buffer (PBS with 0.1% BSA and 2 mM EDTA). Samples were filtered through a 50 μm nylon filter mesh to remove clumps prior to analysis with a five-laser LSR Fortessa (BD Bioscience). Acquired data were analyzed using FlowJo 10.7.2.

## Author Contributions

G.G., R.H.S., and A.D. designed experiments and prepared the manuscript. G.G. performed all *C. elegans* experiments. S.M. and R.H.S. assisted in *C. elegans* experiments. C.K.T. and B.W. designed and created all cell lines. H.Z. performed all cell culture experiments. H.Z. and B.W. assisted in cell culture data analysis. H.Z., C.K.T., and R.H.S. assisted in manuscript preparation.

## Data Availability

All data required to evaluate the conclusions in this manuscript are available within the manuscript and supplementary data. All strains synthesized in this manuscript are derivatives of strains from CGC and are available upon request.

## Acknowledgements

We are grateful to all the members of the Dillin lab for intellectual and technical support. We are grateful to the Dernburg lab for use of their Marianas spinning-disc confocal platform. This work was supported by the following grants: G.G. is supported by T32AG052374, H.Z. is supported by 2020-A-018-FEL through the Larry L. Hillblom Foundation, C.K.T. is supported by F32AG069388 from the National Institute on Aging, R.H.S. is supported by R00AG065200 from the National Institute on Aging, and A.D. is supported by R01AG059566 from the National Institute on Aging and the Howard Hughes Medical Institute. Some strains were provided by the CGC, which is funded by the NIH Office of Research Infrastructure Programs (P40 OD010440).

---

**Table S1.** Candidate LD proteins identified by proteome meta-analysis and screen score. Proteins identified in meta-analysis of published LD isolation proteomes [31]–[33]. Gene description, screen score (scored ± 3 in 0.5 increments in comparison to Empty Vector/*sec-11* RNAi control), and approximated developmental stage at time of screen noted. N/A corresponds to genes not screened due to RNAi availability.

| Gene Name | Sequence Name | Gene Description | Screen Score | Screen Larval Stage |
| --- | --- | --- | --- | --- |
| inf-1 | F57B9.6 | Eukaryotic initiation factor 4A | -3 | L4/Adult |
| rpl-10 | F10B5.1 | 60S ribosomal protein L10 | -3 | L4/Adult |
| rpl-13 | C32E8.2 | 60S ribosomal protein L13 | -3 | L3/L4 |
| rpl-16 | M01F1.2 | 60S ribosomal protein L13a | -3 | L2/L3 |
| rpl-17 | Y48G8AL.8 | 60S ribosomal protein L17 | -3 | L4/Adult |
| rpl-19 | C09D4.5 | 60S ribosomal protein L19 | -3 | L3 |
| rpl-20 | E04A4.8 | 60S ribosomal protein L18a | -3 | Adult |
| rpl-23 | B0336.10 | 60S ribosomal protein L23 | -3 | Adult |
| rpl-3 | F13B10.2 | 60S ribosomal protein L3 | -3 | L3 |
| rpl-35 | ZK652.4 | 60S ribosomal protein L35 | -3 | L3/L4 |
| rpl-9 | R13A5.8 | 60S ribosomal protein L9 | -3 | L4/Adult |
| rps-0 | B0393.1 | 40S ribosomal protein SA | -3 | Adult |
| rps-1 | F56F3.5 | 40S ribosomal protein S3a | -3 | L3 |
| rps-10 | D1007.6 | Ribosomal Protein, Small subunit | -3 | Adult |
| rps-16 | T01C3.6 | 40S ribosomal protein S16 | -3 | Adult |
| rps-18 | Y57G11C.16 | Ribosomal Protein, Small subunit | -3 | L3/L4 |
| rps-19 | T05F1.3 | 40S ribosomal protein S19 | -3 | Adult |
| rps-20 | Y105E8A.16 | Ribosomal Protein, Small subunit | -3 | L4/Adult |
| rps-3 | C23G10.3 | 40S ribosomal protein S3 | -3 | L3/L4 |
| rps-5 | T05E11.1 | 40S ribosomal protein S5 | -3 | Adult |
| rps-8 | F42C5.8 | 40S ribosomal protein S8 | -3 | L4/Adult |
| rps-9 | F40F8.10 | 40S ribosomal protein S9 | -3 | L3/L4 |
| atp-1 | H28O16.1 | ATP synthase subunit alpha, mitochondrial | -2.5 | L3/L4 |
| eef-1A.1 | F31E3.5 | Elongation factor 1-alpha | -2.5 | Adult |
| eef-2 | F25H5.4 | Elongation factor 2 | -2.5 | Adult |
| rla-0 | F25H2.10 | 60S acidic ribosomal protein P0 | -2.5 | Adult |
| rpl-2 | B0250.1 | 60S ribosomal protein L8 | -2.5 | L4/Adult |
| rpl-22 | C27A2.2 | 60S ribosomal protein L22 | -2.5 | L4/Adult |
| rpl-27 | C53H9.1 | 60S ribosomal protein L27 | -2.5 | Adult |
| rpl-36 | F37C12.4 | 60S ribosomal protein L36 | -2.5 | L4/Adult |
| rpl-6 | R151.3 | 60S ribosomal protein L6 | -2.5 | Adult |
| rps-14 | F37C12.9 | 40S ribosomal protein S14 | -2.5 | Adult |
| rps-15 | F36A2.6 | 40S ribosomal protein S15 | -2.5 | Adult |
| rps-22 | F53A3.3 | Ribosomal Protein, Small subunit | -2.5 | L4/Adult |
| rps-7 | ZC434.2 | 40S ribosomal protein S7 | -2.5 | Adult |
| atp-2 | C34E10.6 | ATP synthase subunit beta, mitochondrial | -2 | L3/L4 |
| fib-1 | T01C3.7 | rRNA 2'-O-methyltransferase fibrillarin | -2 | Adult |
| rab-1 | C39F7.4 | RAB family | -2 | L3/L4 |
| rpl-18 | Y45F10D.12 | 60S ribosomal protein L18 | -2 | Adult |
| rpl-5 | F54C9.5 | 60S ribosomal protein L5 | -2 | Adult |
| rps-11 | F40F11.1 | Ribosomal Protein, Small subunit | -2 | Adult |
| rps-2 | C49H3.11 | 40S ribosomal protein S2 | -2 | Adult |
| hsp-1 | F26D10.3 | Heat shock 70 kDa protein A | -1.5 | Adult |
| hsp-90 | C47E8.5 | Heat shock protein 90 | -1.5 | Adult |
| let-767 | C56G2.6 | Very-long-chain 3-oxoacyl-coA reductase let-767 | -1.5 | Adult |
| tba-2 | C47B2.3 | Tubulin alpha-2 chain | -1.5 | Adult |
| ucr-1 | F56D2.1 | Cytochrome b-c1 complex subunit 1, mitochondrial | -1.5 | Adult |
| vha-13 | Y49A3A.2 | V-type proton ATPase catalytic subunit A | -1.5 | L3/L4 |
| vha-15 | T14F9.1 | Probable V-type proton ATPase subunit H 2 | -1.5 | L3/L4 |
| act-4 | M03F4.2 | Actin-4 | -1 | Adult |
| ant-1.1 | T27E9.1 | Adenine Nucleotide Translocator | -1 | Adult |
| atad-3 | F54B3.3 | ATPase family AAA domain-containing protein 3 | -1 | Adult |
| eat-6 | B0365.3 | Sodium/potassium-transporting ATPase subunit alpha | -1 | Adult |
| eef-1G | F17C11.9 | Probable elongation factor 1-gamma | -1 | L4/Adult |
| F42G8.10 | F42G8.10 | NADH:ubiquinone oxidoreductase subunit B11 | -1 | Adult |
| hsp-60 | Y22D7AL.5 | Chaperonin homolog Hsp-60, mitochondrial | -1 | Adult |
| T02H6.11 | T02H6.11 | ubiquinol-cytochrome c reductase binding protein | -1 | Adult |
| tba-4 | F44F4.11 | Tubulin alpha chain | -1 | Adult |
| Y7A5A.1 | Y7A5A.1 | 24-dehydrocholesterol reductase | -1 | Adult |
| acs-17 | C46F4.2 | Fatty Acid CoA Synthetase family | -0.5 | Adult |
| ant-1.3 | K01H12.2 | Adenine Nucleotide Translocator; Mitochondrial | -0.5 | Adult |
| B0491.5 | B0491.5 | enriched in the OLL, the PVD, and the germ line | -0.5 | Adult |
| gdh-1 | ZK829.4 | Glutamate dehydrogenase | -0.5 | Adult |
| ldp-1 | F22F7.1 | Lipid droplet localized protein | -0.5 | Adult |
| nuo-2 | T10E9.7 | NADH Ubiquinone Oxidoreductase | -0.5 | Adult |
| rab-7 | W03C9.3 | RAB family | -0.5 | Adult |
| rack-1 | K04D7.1 | Guanine nucleotide-binding protein subunit beta-2-like 1 | -0.5 | Adult |
| rpl-31 | W09C5.6 | 60S ribosomal protein L31 | -0.5 | Adult |
| rps-4 | Y43B11AR.4 | 40S ribosomal protein S4 | -0.5 | Adult |
| algn-2 | F09E5.2 | Asparagine Linked Glycosylation (ALG) homolog, Nematode | 0 | Adult |
| alh-8 | F13D12.4 | Probable methylmalonate-semialdehyde dehydrogenase | 0 | Adult |
| cgh-1 | C07H6.5 | ATP-dependent RNA helicase cgh-1 | 0 | Adult |
| cts-1 | T20G5.2 | Probable citrate synthase, mitochondrial | 0 | Adult |
| D2030.4 | D2030.4 | NADH dehydrogenase | 0 | Adult |
| dhs-19 | T11F9.11 | DeHydrogenases, Short chain | 0 | Adult |
| dhs-3 | T02E1.5 | DeHydrogenases, Short chain | 0 | Adult |
| F01G4.6 | F01G4.6 | Phosphate carrier protein, mitochondrial | 0 | Adult |
| F44B9.5 | F44B9.5 | Ancient ubiquitous protein 1 homolog | 0 | Adult |
| F52H2.6 | F52H2.6 | Diminuto-like protein | 0 | Adult |
| lec-1 | W09H1.6 | 32 kDa beta-galactoside-binding lectin | 0 | Adult |
| par-5 | M117.2 | 14-3-3-like protein 1 | 0 | Adult |
| pdi-2 | C07A12.4 | Protein disulfide-isomerase 2 | 0 | Adult |
| phb-2 | T24H7.1 | Mitochondrial prohibitin complex protein 2 | 0 | Adult |
| plin-1 | W01A8.1 | Perilipin-1 homolog | 0 | Adult |
| rpl-32 | T24B8.1 | Ribosomal Protein, Large subunit | 0 | Adult |
| tba-7 | T28D6.2 | Tubulin alpha chain | 0 | Adult |
| tomm-20 | F23H12.2 | Mitochondrial import receptor subunit TOM20 homolog | 0 | Adult |
| unc-108 | F53F10.4 | RAB2A, member RAS oncogene family | 0 | Adult |
| vdac-1 | R05G6.7 | Probable voltage-dependent anion-selective channel | 0 | Adult |
| vit-1 | K09F5.2 | Vitellogenin-1 | 0 | Adult |
| vit-2 | C42D8.2 | Vitellogenin-2 | 0 | Adult |
| vit-6 | K07H8.6 | Vitellogenin-6 | 0 | Adult |
| acs-4 | F37C12.7 | Fatty Acid CoA Synthetase family | 0.5 | Adult |
| tbb-1 | K01G5.7 | Tubulin beta chain | 0.5 | Adult |
| alh-4 | T05H4.13 | Aldehyde dehydrogenase | 1 | Adult |
| enpl-1 | T05E11.3 | Endoplasmin homolog | 1 | Adult |
| hsp-3 | C15H9.6 | Heat shock 70 kDa protein C | 1 | Adult |
| vit-5 | C04F6.1 | Vitellogenin-5 | 1 | Adult |
| hsp-4 | F43E2.8 | Endoplasmic reticulum chaperone BiP homolog | 2 | Adult |
| ben-1 | C54C6.2 | Tubulin beta chain | N/A | N/A |
| dhs-9 | Y32H12A.3 | DeHydrogenases, Short chain | N/A | N/A |
| eef-1A.2 | R03G5.1 | Elongation factor 1-alpha | N/A | N/A |
| faah-4 | Y56A3A.12 | Fatty Acid Amide Hydrolase homolog | N/A | N/A |
| rpl-1 | Y71F9AL.13 | 60S ribosomal protein L10a | N/A | N/A |
| rpl-15 | K11H12.2 | 60S ribosomal protein L15 | N/A | N/A |
| rpl-21 | C14B9.7 | 60S ribosomal protein L21 | N/A | N/A |
| rpl-4 | B0041.4 | 60S ribosomal protein L4 | N/A | N/A |
| rpl-7 | F53G12.10 | 60S ribosomal protein L7 | N/A | N/A |
| rpl-7A | Y24D9A.4 | 60S ribosomal protein L7a | N/A | N/A |
| rps-13 | C16A3.9 | 40S ribosomal protein S13 | N/A | N/A |
| rps-28 | Y41D4B.5 | 40S ribosomal protein S28 | N/A | N/A |
| rps-6 | Y71A12B.1 | 40S ribosomal protein S6 | N/A | N/A |
| sip-1 | F43D9.4 | Stress-induced protein 1 | N/A | N/A |
| trap-1 | Y71F9AM.6 | TRanslocon-Associated Protein | N/A | N/A |
| tsn-1 | F10G7.2 | Staphylococcal nuclease domain-containing protein | N/A | N/A |
| vha-11 | Y38F2AL.3 | V-type proton ATPase subunit C | N/A | N/A |
| vit-4 | F59D8.2 | Vitellogenin-4 | N/A | N/A |
| Y53F4B.18 | Y53F4B.18 | fatty acid amide hydrolase 2 | N/A | N/A |
| Y69A2AR.18 | Y69A2AR.18 | ATP synthase F1 subunit gamma | N/A | N/A |
| ZK809.3 | ZK809.3 | X-ray repair cross complementing 5 | N/A | N/A |

**Table S2.** Statistical analysis of tunicamycin survival assay data. Median lifespan, death events counted, and statistics for tunicamycin survival assay of worms grown on Empty Vector (EV) or *let-767* RNAi combined with EV or *sbp-1* RNAi.

| Supplemental Table S2. Statistical Analyses of Tunicamycin Survival Data |  |  |  |  |  |
| --- | --- | --- | --- | --- | --- |
| Corresponding | Strain, Treatment | Median Lifespan | # Deaths/# Total | % Change in | p-value |
| Figure |  | [days] |  | Median Lifespan | Log-rank (Mantel-Cox) |
| 3E | N2, Empty Vector RNAi, 25ng/μL tunicamycin | 14 | 71/100 | n/a | n/a |
|  | N2, *sbp-1*/Empty Vector RNAi, 25ng/μL tunicamycin | 15 | 97/100 | 7.1 | 0.6013 |
|  | N2, *let-767*/Empty Vector RNAi, 25ng/μL tunicamycin | 8 | 67/100 | -42.9 | <0.0001 |
|  | N2, *let-767/sbp-1* RNAi, 25ng/μL tunicamycin | 14 | 97/100 | 0.0 | 0.3637 |
| 3F | N2, vector RNAi, 1% DMSO | 21 | 76/100 | n/a | n/a |
|  | N2, *sbp-1*/Empty Vector RNAi, 1% DMSO | 14 | 96/100 | -33.3 | <0.0001 |
|  | N2, *let-767*/Empty Vector RNAi, 1% DMSO | 8 | 33/50 | -61.9 | <0.0001 |
|  | N2, *let-767/sbp-1* RNAi, 1% DMSO | 13 | 96/13 | -38.1 | <0.0001 |
| Replicates |  |  |  |  |  |
| 3E/F V2 | N2, Empty Vector RNAi, 25ng/μL tunicamycin | 12 | 71/100 | n/a | n/a |
|  | N2, *sbp-1*/Empty Vector RNAi, 25ng/μL tunicamycin | 14 | 85/100 | 16.7 | 0.7 |
|  | N2, *let-767*/Empty Vector RNAi, 25ng/μL tunicamycin | 8 | 42/100 | -33.3 | <0.0001 |
|  | N2, *let-767/sbp-1* RNAi, 25ng/μL tunicamycin | 14 | 86/100 | 16.7 | 0.7 |
|  | N2, vector RNAi, 1% DMSO | 19 | 66/100 | n/a | n/a |
|  | N2, *sbp-1*/Empty Vector RNAi, 1% DMSO | 15 | 84/100 | -21.1 | <0.0001 |
|  | N2, *let-767*/Empty Vector RNAi, 1% DMSO | 7 | 29/100 | -63.2 | <0.0001 |
|  | N2, *let-767/sbp-1* RNAi, 1% DMSO | 15 | 83/100 | -21.1 | <0.0001 |
| 3E/F V3 | N2, Empty Vector RNAi, 25ng/μL tunicamycin | 11 | 114/120 | n/a | n/a |
|  | N2, *sbp-1*/Empty Vector RNAi, 25ng/μL tunicamycin | 11 | 115/120 | 0.0 | 0.7 |
|  | N2, *let-767*/Empty Vector RNAi, 25ng/μL tunicamycin | 6 | 102/120 | -45.5 | <0.0001 |
|  | N2, *let-767/sbp-1* RNAi, 25ng/μL tunicamycin | 11 | 114/121 | 0.0 | 0.7 |
|  | N2, vector RNAi, 1% DMSO | 17 | 102/120 | n/a | n/a |
|  | N2, *sbp-1*/Empty Vector RNAi, 1% DMSO | 13 | 100/120 | -23.5 | <0.0001 |
|  | N2, *let-767*/Empty Vector RNAi, 1% DMSO | 6 | 47/120 | -64.7 | <0.0001 |
|  | N2, *let-767/sbp-1* RNAi, 1% DMSO | 12 | 98/120 | -29.4 | <0.0001 |

**Figure S1.**
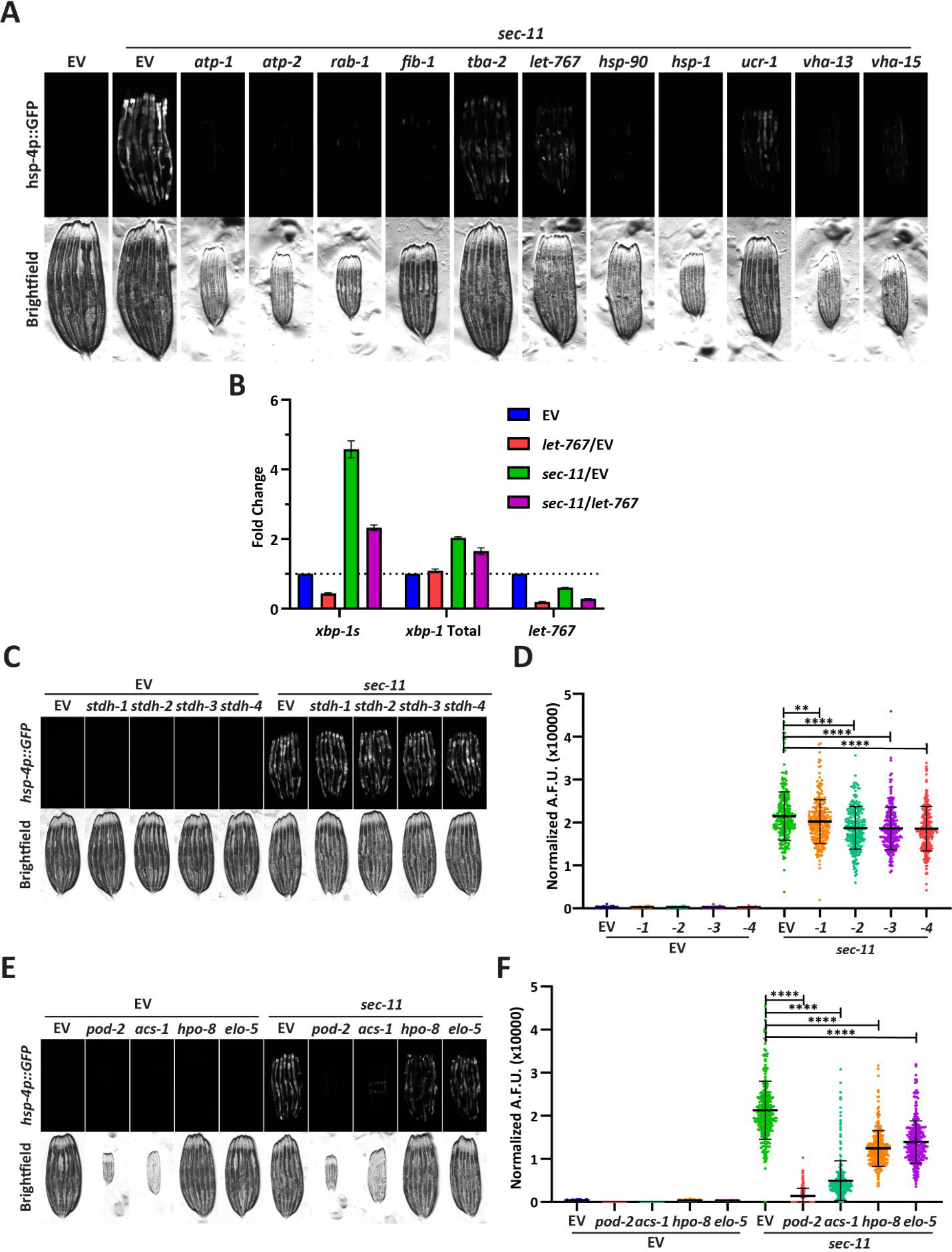
Knockdown of *let-767* suppresses the UPR^ER^ more severely than other lipid related genes. (A) Fluorescent micrographs of transgenic animals expressing *hsp-4p∷GFP* grown from L1 on Empty Vector (EV) or *sec-11* RNAi combined in a 1:1 ratio with either *atp-1*, *atp-2*, *rab-1*, *fib-1*, *tba-2*, *let-767*, *hsp-90*, *hsp-1*, *ucr-1*, *vha-13*, or *vha-15* RNAi and imaged at day 1 of adulthood to assay effects on UPR^ER^ induction. (B) Quantitative RT-PCR transcript levels of *xbp-1s,* total, *xbp-1*, and *let-767* from day 1 adult N2 animals grown from L1 on EV or *sec-11* RNAi combined in a 1:1 ratio with either EV or *let-767* RNAi. Fold-change compared to EV treated N2 animals. Error bars indicate ± standard deviation across 3 biological replicates, each averaged from 2 technical replicates. (C) Fluorescent micrographs of day 1 adult transgenic animals expressing *hsp-4p∷GFP* grown from L1 on EV, *stdh-1*, *stdh-2*, *stdh-3*, or *stdh-4* RNAi combined in a 1:1 ratio with either EV or *sec-11* RNAi to assay effects on UPR^ER^ induction. (D) Quantification of (C) using a BioSorter. Lines represent mean and standard deviation. n = 245. Mann-Whitney test P-value **< 0.05, **** < 0.0001. Data is representative of 3 replicates. (E) Fluorescent micrographs of transgenic animals expressing *hsp-4p∷GFP* grown from L1 on EV, *pod-2*, *acs-1*, *hpo-8*, or *elo-5* RNAi combined in a 1:1 ratio with either EV or *sec-11* RNAi to assay effects on UPR^ER^ induction. (F) Quantification of (E) using a BioSorter. Lines represent mean and standard deviation. n = 236. Mann-Whitney test P-value **** < 0.0001. Data is representative of 2 replicates.

**Figure S2.**
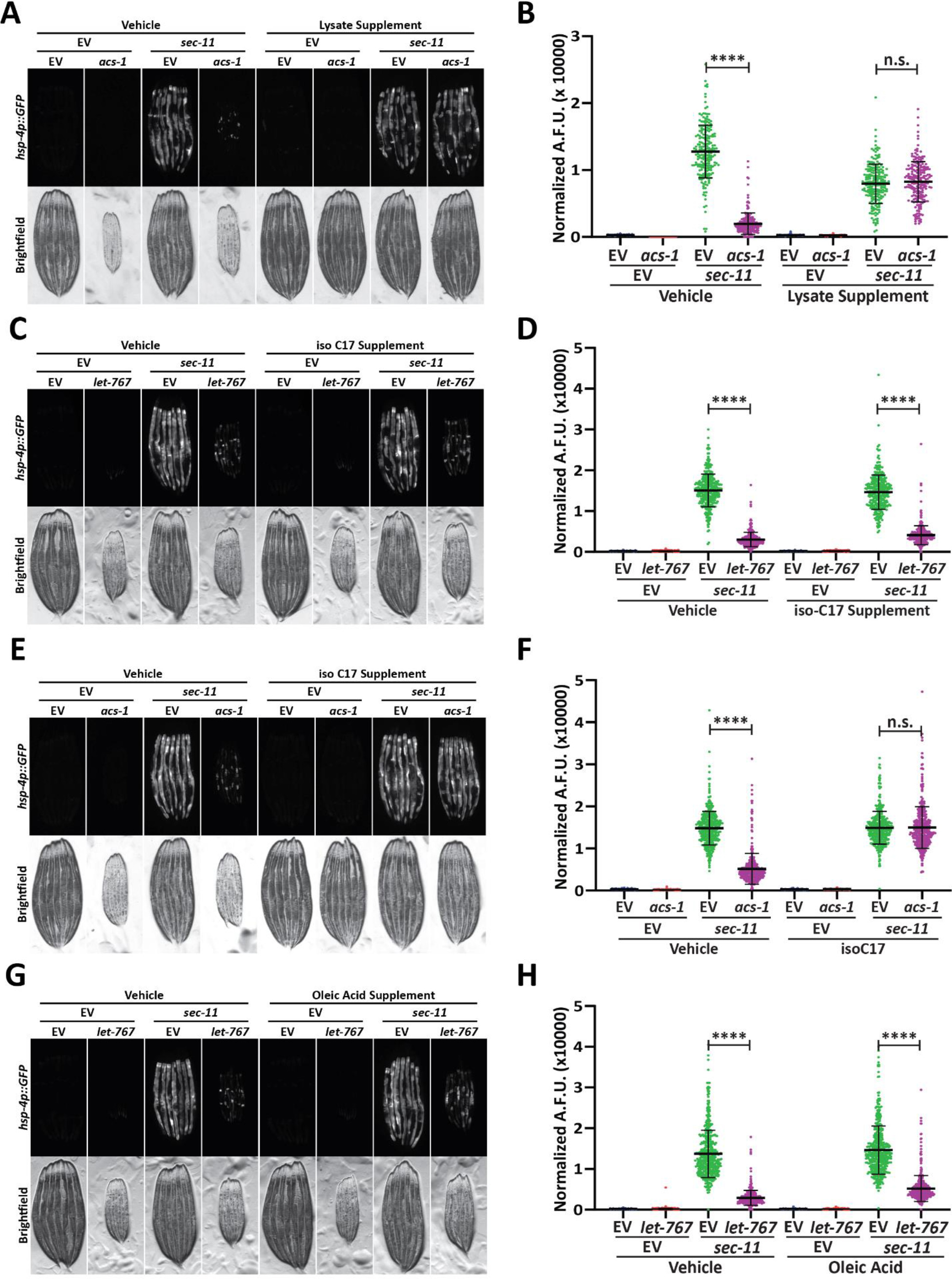
Supplementation of mmBCFA or oleic acid does not restore the UPR^ER^ suppressed by *let-767* RNAi. (A) Fluorescent micrographs of transgenic animals expressing *hsp-4p∷GFP* grown on EV or *acs-1* RNAi combined in a 1:1 ratio with either EV or *sec-11* RNAi supplemented with vehicle or N2 lysate to assay supplementation efficacy. (B) Quantification of (A) using a Biosorter. Lines represent mean and standard deviation. n = 211. Mann-Whitney test P-value **** < 0.0001. n.s. = non-significant. Data is representative of 3 replicates. (C) Fluorescent micrographs of day 1 adult transgenic animals expressing *hsp-4p∷GFP* grown on Empty Vector (EV) or *let-767* RNAi combined in a 1:1 ratio with either EV or *sec-11* RNAi grown with vehicle or 0.5mM isoC17 to assay effects on UPR^ER^ induction. (D) Quantification of (C) using a BioSorter. Lines represent mean and standard deviation. n = 338. Mann-Whitney test P-value **** < 0.0001. Data is representative of 3 replicates. (E) Fluorescent micrographs of day 1 adult transgenic animals expressing *hsp-4p∷GFP* grown on EV or *acs-1* RNAi combined in a 1:1 ratio with either EV or *sec-11* RNAi grown with vehicle or 0.5mM isoC17 to assay effects on UPR^ER^ induction. (F) Quantification of (E) using a BioSorter. Lines represent mean and standard deviation. n = 428. Mann-Whitney test P-value **** < 0.0001, n.s. = non-significant. Data is representative of 3 replicates. (G) Fluorescent micrographs of day 1 adult transgenic animals expressing *hsp-4p∷GFP* grown on EV or *let-767* RNAi combined in a 1:1 ratio with either EV or *sec-11* RNAi grown with vehicle or 1mM oleic acid to assay effects on the UPR^ER^ induction. (H) Quantification of (G) using a BioSorter. Lines represent mean and standard deviation. n = 461. Mann-Whitney test P-value **** < 0.0001. Data is representative of 3 replicates.

**Figure S3.**
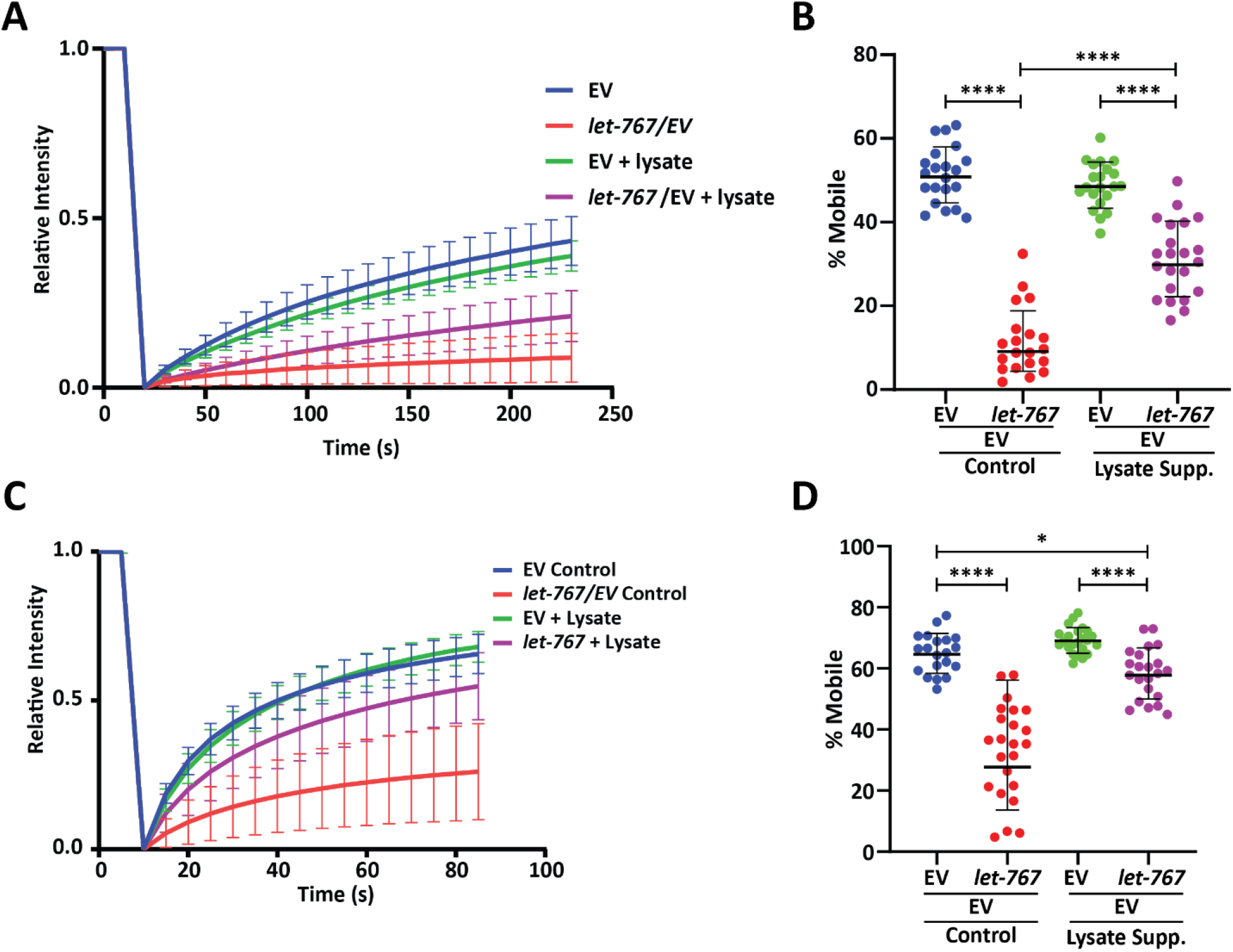
Lysate supplementation rescues luminal ER protein dynamics but not ER membrane protein dynamics. (A) FRAP curve of intestinal ER-transmembrane protein, SPCS-1∷GFP, from day 1 adult animals grown on Empty Vector (EV) or 50% *let-767* RNAi supplemented with vehicle or N2 lysate (geometric mean for n = 20 pooled from 2 biological replicates). Lines represent geometric standard deviation. (B) Calculated percent mobile SPCS-1∷GFP of (A). Lines represent geometric mean and geometric standard deviation. Mann-Whitney test P-value **** < 0.0001. (C) FRAP curve of intestinal ER-lumen protein, mRuby∷HDEL, from day 1 adult animals grown on Empty Vector (EV) or 50% *let-767* RNAi supplemented with vehicle or N2 lysate (geometric mean for n = 20 pooled from 2 biological replicates). Lines represent geometric standard deviation. (D) Calculated percent mobile mRuby∷HDEL of (A). Lines represent geometric mean and geometric standard deviation. Mann-Whitney test P-value **** < 0.0001 and * < 0.05.

**Figure S4.**
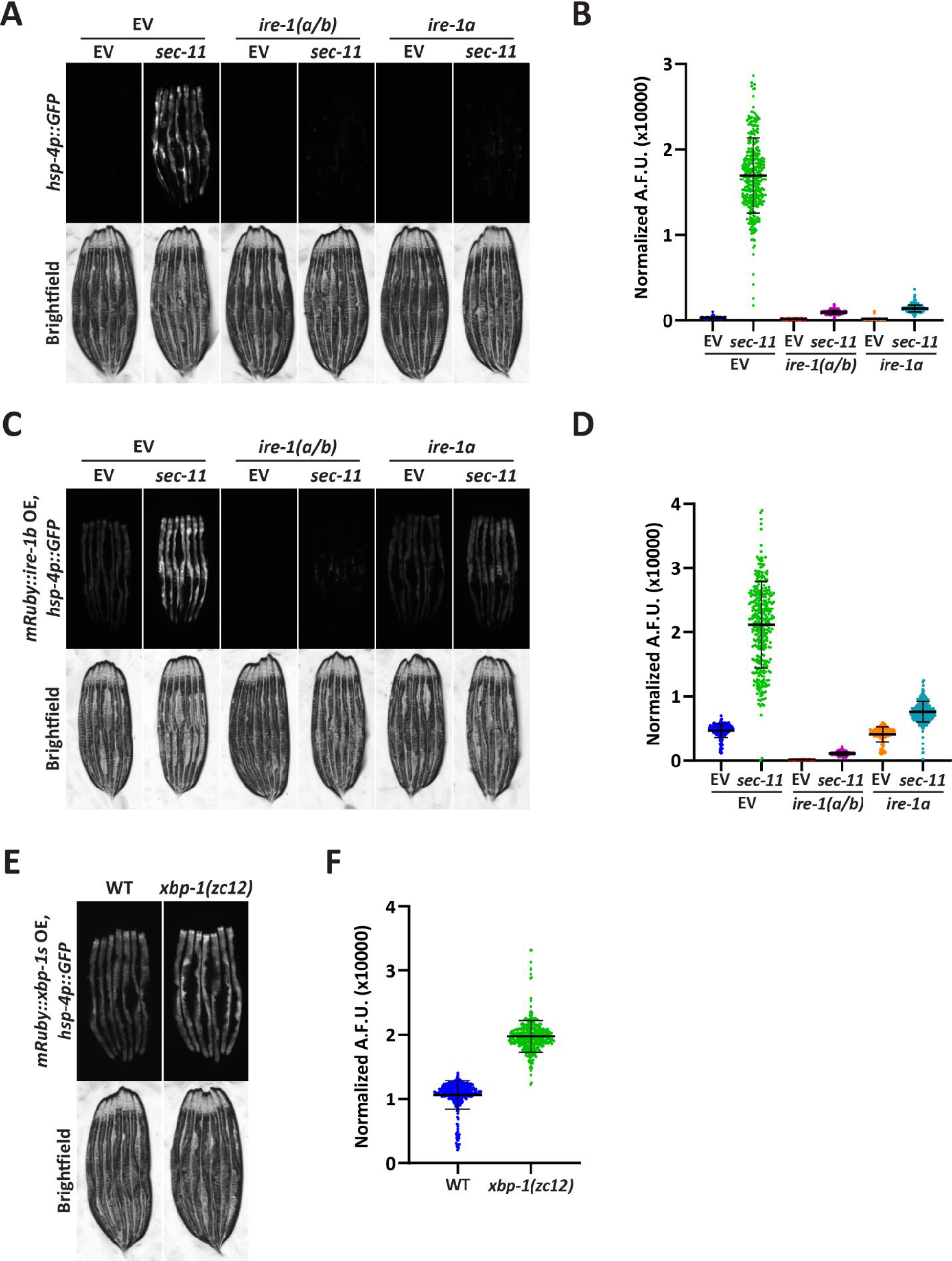
Overexpression of *ire-1b* or *xbp-1s* induces the UPR^ER^ independently of endogenous *ire-1a* or *xbp-1*, respectively. (A) Fluorescent micrographs of transgenic animals expressing *hsp-4p∷GFP* grown on Empty Vector (EV), *ire-1*, or *ire-1a* RNAi combined with either EV or *sec-11* RNAi to assay the UPR^ER^ induction. (B) Quantification of (A) using a BioSorter. Lines represent mean and standard deviation. n = 321. Mann-Whitney test P-value **** < 0.0001. Data is representative of 2 replicates. (C) Fluorescent micrographs of day 1 adult transgenic animals expressing *hsp-4p∷GFP* and intestinal *mRuby∷ire-1b* grown on EV, *ire-1*, or *ire-1a* RNAi combined with either EV or *sec-11* RNAi to assay the UPR^ER^ induction. (D) Quantification of (C) using a BioSorter. Lines represent mean and standard deviation. n = 344. Mann-Whitney test P-value **** < 0.0001. Data is representative of 2 replicates. (E) Fluorescent micrographs of day 1 adult transgenic animals expressing *hsp-4p∷GFP* and intestinal *mRuby∷xbp-1s* in a WT or *xbp-1 (zc12)* null-mutant background grown on EV to assay the UPR^ER^ induction. (F) Quantification of (E) using a BioSorter. Lines represent mean and standard deviation. n = 367. Mann-Whitney test P-value **** < 0.0001. Data is representative of 2 replicates.

